# Differences in Pneumococcal Serotype Replacement in Individuals with and without Underlying Medical Conditions

**DOI:** 10.1101/296772

**Authors:** Daniel M. Weinberger, Joshua L Warren, Tine Dalby, Eugene D. Shapiro, Valentiner Branth, Hans-Christian Slotved, Zitta Barrella Harboe

## Abstract

**Background:** Pneumococcal conjugate vaccines (PCVs) have had a well-documented impact on the incidence of invasive pneumococcal disease (IPD) worldwide. However, declines in IPD due to vaccine-targeted serotypes have been partially offset by increases in IPD due to non-vaccine serotypes. The goal of this study was to quantify serotype-specific changes in the incidence of IPD that occurred in different age groups, with or without certain co-morbidities, following the introduction of PCV7 and PCV13 in the childhood vaccination program in Denmark.

**Methods:** We used nationwide surveillance data for IPD in Denmark and a hierarchical Bayesian regression framework to estimate changes in the incidence of IPD associated with the introduction of PCV7 (2007) and PCV13 (2010) while controlling for serotype-specific epidemic cycles and unrelated secular trends.

**Results and Conclusions:** Following the introduction of PCV7 and 13 in children, the net impact of serotype replacement varied considerably by age group and the presence of comorbid conditions. Serotype replacement offset a greater fraction of the decline in vaccine-targeted serotypes following the introduction of PCV7 compared with the period following the introduction of PCV13. Differences in the magnitude of serotype replacement were due to variations in the incidence of non-vaccine serotypes in the different risk groups before the introduction of PCV7 and PCV13. The relative increases in the incidence of IPD caused by non-vaccine serotypes did not differ appreciably in the post-vaccination period. Serotype replacement offset a greater proportion of the benefit of PCVs in strata in which the non-vaccine serotypes comprised a larger proportion of cases prior to the introduction of the vaccines. These findings could help to predict the impact of next-generation conjugate vaccines in specific risk groups.

## RESEARCH IN CONTEXT

### Evidence before this study

We searched PubMed for studies of pneumococcal vaccine impact from 2000 to the present using the search term “pneumococ*”. Following the introduction of the original pneumococcal conjugate vaccine (PCV7), the serotypes targeted by the vaccine were nearly eliminated as causes of invasive pneumococcal disease (IPD), a severe form of pneumococcal infection. These reductions were seen among both young children (who received the vaccine) and among adults (who did not receive the vaccine but indirectly benefit from a reduction in transmission of the bacteria). Despite these reductions, serotypes not targeted by the vaccine increased as causes of IPD, partially reducing the benefit of the vaccine. Following the switch to PCV13, fewer data are available, but there again were strong declines in IPD due to vaccine-targeted serotypes and increases in non-targeted serotypes. Cross-country comparisons of the impacts of PCV7 demonstrated that the net benefit of the vaccine varied considerably between populations, and in some countries and some age groups, serotype replacement completely offset the reduction IPD due to vaccine-targeted serotypes. The proportion of healthy children carrying vaccine-targeted serotypes in the population prior to vaccine introduction partially explains the differences between countries. However, the impact of serotype replacement on the net reduction in disease rates also varies between age groups and those with and without comorbidity, and these variations are not as well understood.

### Added value of this study

In this study, we describe how declines in vaccine-targeted serotypes and increases in non-vaccine serotypes have affected individuals of different age groups, with and without underlying comorbidities. To do this, we used high-quality data from nationwide health registers and surveillance data from Denmark. The net benefit of PCV7 and PCV13 varied considerably by age group and by presence of comorbidity. Following PCV7, serotype replacement completely offset declines in vaccine-targeted serotypes in adults with and without co-morbidities. However, following PCV13, all risk groups exhibited substantial declines in the incidence of invasive pneumococcal disease. Serotype replacement offset a greater proportion of the benefit in risk groups in which non-vaccine serotypes comprised a larger proportion of cases prior to the introduction of the vaccine. However, the relative increase in individual non-vaccine serotypes was similar between risk groups. This provides a novel understanding of the variations in the impact of PCVs among different risk groups living in the same population. This information could help to predict the impact of next-generation conjugate vaccines in specific risk groups.

### Implications of all the available evidence

Despite clear evidence of serotype replacement, the use of PCV13 in infants has a net benefit in preventing severe disease in all age groups, with and without comorbidity. However, the net benefit of pneumococcal conjugate vaccines varies by age and presence of comorbidity. Next-generation conjugate vaccines that target additional serotypes should focus on these emerging serotypes that disproportionately affect specific risk groups.

## INTRODUCTION

Pneumococcal conjugate vaccines (PCVs) have had a well-documented impact on the incidence of invasive pneumococcal disease (IPD) worldwide.^1-5^ Because PCVs interrupt transmission of pneumococcus among healthy children (the major reservoir for the bacteria), the introduction of PCVs has led to reductions in the incidence of IPD among both vaccinated children and (indirectly) among unvaccinated adults (herd protection).^6,7^ A meta-analysis of data from the United States, Europe, and Australia found that on average, IPD due to serotypes in PCV7 declined by ~90% among adults ^1^. However, these declines in IPD due to vaccine-targeted serotypes were at least partially offset by increases in IPD caused by non-vaccine serotypes, a phenomenon known as “serotype replacement.^1,8^ The net benefit of PCVs is therefore determined by both the magnitude of the decline in the incidence of IPD due to serotypes targeted by PCVs and by increases in the incidence of serotypes not targeted by the vaccine. The net benefit of PCVs can vary by risk groups and between populations.^1,9^ Variations in the net benefit of PCVs between countries can be explained by variations in the serotype distribution of nasopharyngeal carriage among children in the pre-vaccine period.^10^ However, it is not clear why the net benefit of PCVs vary within the same population where exposure is more consistent between risk groups. Understanding variations in the net benefit of PCVs against IPD in different risk groups is important as conjugate vaccines with expanded numbers of serotypes are developed and deployed.^11^

Denmark is an ideal setting for evaluating this issue. The 7-valent vaccine (PCV7) was introduced in the childhood immunization program in 2007 and was replaced in 2010 with the 13-valent vaccine (PCV13). Nationwide surveillance for IPD has been conducted consistently for decades, and a nationwide health registry allows for the identification of underlying conditions in individuals who develop IPD.

The goal of this study was to quantify and explain changes in the incidence of IPD caused by vaccine and non-vaccine serotypes after the introduction of PCVs in children among different groups of patients in Denmark according to their age and presence of underlying diseases.

Distinguishing changes in the incidence of IPD due to the effects by PCVs from those due to unrelated secular trends is a major challenge because of the well-recognized multi-year, serotype-specific epidemic cycles.^12,13^ To do this, we used an analytical framework that estimates and controls for the epidemic cycles of individual serotypes and can estimate serotype- and risk– group-specific estimates of vaccine effects against IPD as well as average estimates across all serotypes and risk groups.

## METHODS

### Data sources

Data were retrieved by linking several Danish national registers from 1994-2015. The data on cases of IPD (i.e., meningitis, bacteremia, and other sterile foci) were obtained from a laboratory based nationwide surveillance system that captures more than 95% of all laboratory-confirmed IPD-cases in Denmark. Reporting in this system has been consistent since the early 1990s. The serotype is determined for more than 97% of all IPD isolates at the Statens Serum Institut. This study was approved by the Danish Data Protection Agency (2012-41-1068).

PCV7 was introduced in the Danish childhood immunization program in 2007, and then replaced by PCV13 in 2010. PCVs have been administered free of charge in a 2+1 schedule at the age of 3,5, and 12 months. The uptake of the vaccine is high at more than 90% for the full series and similar to the uptake of the pentavalent vaccine (Diphteria, Tetanus, acellular pertussis, IVP, capsular *Haemophilus influenzae*) given concomitantly in the program. Throughout the manuscript, ‘PCV7 serotypes’ refer to serotypes 4 6B, 9V, 14, 18C, 19F, and 23F. ‘PCV13 serotypes’ include PCV7 serotypes plus 1, 3, 5, 6A, 7F, and 19A. All other serotypes were classified as non-PCV7/13 serotypes. Prior to 2006/07, serotype 6C isolates were not identifiable and would have been mis-classified as serotype 6A.^14^ The comorbidity status of the cases was ascertained using a record linkage to the Danish National Patient Registry,^15^ a nationwide hospitalization database. The Danish Personal Identification Number, CPR, was used for the linkage. Comorbidity was defined based on discharge codes based on the International Classification of Disease, 10^th^ revision (ICD-10) discharge codes.^16^ ICD-10 codes were included when these had been registered before or at the date of the individual IPD episode. We searched for conditions that increase the risk of IPD as defined by the Advisory Committee on Immunization Practices (ACIP),^17^ with codes related to chronic heart disease, chronic pulmonary disease, diabetes, alcoholism, liver disease, CSF-Leaks, cochlear implants, anatomic or functional asplenia, HIV and other immunodeficiencies, renal disease, hematological and solid malignancies, rheumatological diseases, and organ transplantation. For the specific codes included in the comorbidity analysis, see **Supplementary text**. Information on comorbidities was obtained for all individuals until 2014/15 (ending June 2015). A case was included as an individual belonging to a high-risk group if the patient had a record of any of the indicated conditions at the time of diagnosis with IPD (including if the comorbidity was identified at the time of diagnosis).

In 102 cases (0.5%), we had data on the serogroup but not the serotypes that caused the infection. We assumed that these cases with missing information had the same serotype distribution as the other cases of the same serogroup in the same year and risk group. For instance, if there were 10 cases that were identified as serogroup 6 but without serotype information among adults ≥65 years of age without comorbidity, and the proportion of serotypes 19A, 19F were 0.4 and 0.6 in this strata, respectively, we assumed that among the cases with missing serotype information, 4 were 19A and 6 were 19F.

For estimating the denominator/offset, we obtained the population size in each age group and year from Statistics Denmark^18^. We then calculated the proportion of individuals in each age group who had any of the high-risk conditions using data from up to 19 randomly selected individuals from the background population who did not have IPD. Multiplying the proportion of individuals with comorbid diseases by the population size gave an estimate of the number of individuals in each age group with or without comorbidities.

### Model

There were 74 serotypes included in the analysis, and the data were stratified into five age groups (<5, 5-17, 18-39, 40-64, ≥65 years) and by comorbidity status (with or without known comorbidities prior to the IPD episode). The time series for many of these strata had low counts, making separate model fitting for these strata impractical and the results unreliable.

Moreover, many of the serotypes exhibited multi-year epidemic cycles and trends unrelated to the vaccine that could influence estimates of vaccine-associated changes. To account for these issues, we used a hierarchical Poisson model in the Bayesian setting, where information among different serotypes and risk groups was shared based on the introduction of random effects in the model. We controlled for repeating epidemic cycles and secular trends in each strata. The vaccine-associated declines were captured using linear splines, and we estimated the relative change at each post-vaccine time point as well as the number of cases prevented. The major assumption of this model is that any trends or cycles that occurred during the pre-vaccine period continued in the post-vaccine period. The models were fit using JAGS in R statistical software (https://www.r-project.org/).^19,20^ Details of the model structure, fitting procedures, and calculation of vaccine effects can be found in the Supplementary methods section.

## RESULTS

### Characteristics of data

We included 19,638 cases of IPD that occurred between July 1994 and June 2015 in Denmark. The proportion of cases that were aged ≥65 years increased from 52% in 1994/95 to 64% in 2014/15. Likewise, among the age group ≥65 years the proportion of cases with at least one recorded comorbidity increased steadily from 34% in 1994/95 to 56% in 2000/01 and 74% in 2014/15.

IPD caused by several serotypes had significant multi-year epidemic patterns that were independent of the introduction of PCVs. Epidemic cycles were notable for serotypes 1 (7.7 years; 95%CI: 7.4, 8.1) and 18C (9.7 years, 95% CI: 8.9 years, 10.0 years) (**Figure S5**). The intensity (amplitude) of the IPD epidemic cycles did not differ significantly by risk group. There was evidence of less pronounced harmonic variations in the incidence of IPD due to other serotypes (e.g. 4, 7F) (**Figure S5**).

### Overall changes in IPD associated with the introduction of PCV7 and PCV13

Considering all age and risk groups together, the PCV7-targeted serotypes have declined gradually and have been almost completely eliminated as causes of IPD after the introduction of the vaccine in 2007, declining by 95% between 2007/08 and 2014/15 (Figure 1). The incidence of IPD due to the six additional serotypes in PCV13 increased modestly (36% increase) between 2007 and 2009, along with other non-PCV7 serotypes, and then declined following the introduction of PCV13 in 2010. The incidence of non-vaccine serotype IPD has increased steadily since the introduction of PCV7, having increased by 50% since 2007/08 (Figure 1).

**Figure 1:**
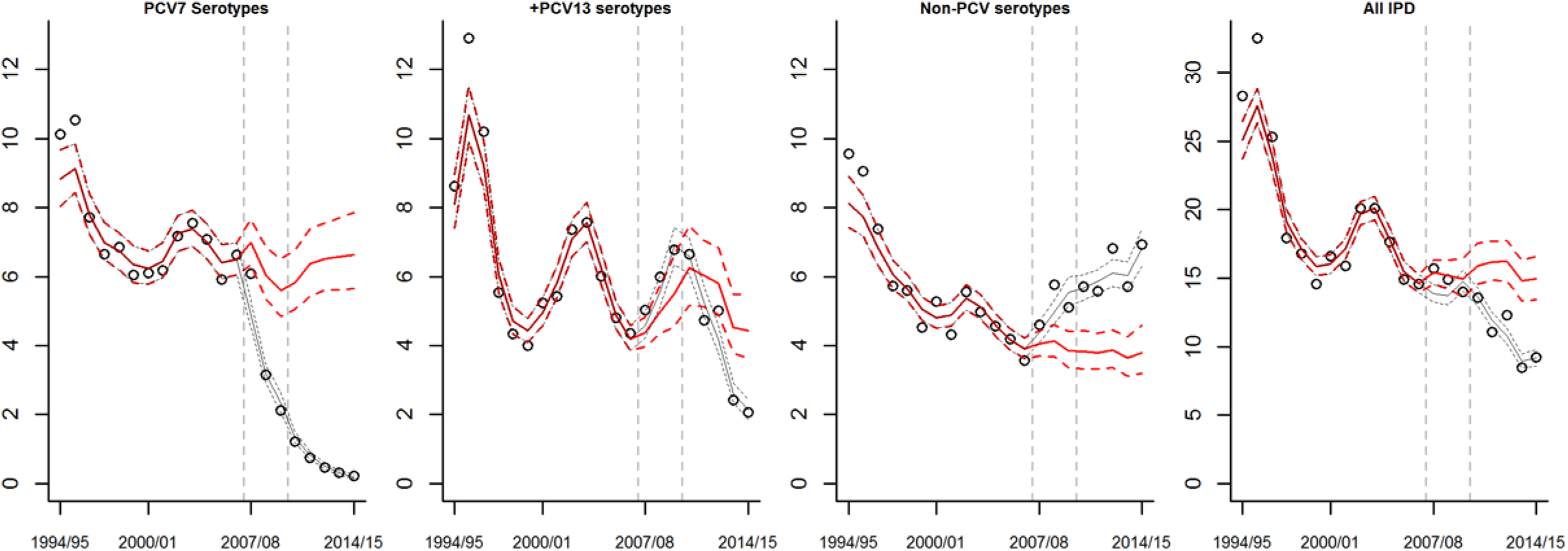
Changes in the incidence of invasive pneumococcal disease (cases/100,000) from 1994/95 to 2014/15 for serotypes targeted by PCV7, for the 6 additional serotypes in PCV13, for non-PCV13 serotypes, and for all cases combined. Incidence is standardized to the age and comorbidity distribution for 2000/01, chosen arbitrarily as a middle point in the time series. The dots represent observed values, the black line+/- 95% credible intervals indicates the model-fitted values, and the red line +/- 95% credible intervals represents the estimate of what the incidence would have been in the absence of vaccine.

These decreases in IPD due to vaccine-targeted serotypes and increases in IPD due to non-vaccine serotypes resulted in a small net-decline in the incidence of IPD between 2007 and 2009 and more substantial declines since the introduction of PCV13 (Table 1). Among individual serotypes, IPD due to serotypes 14 and 19F had the greatest declines in incidence, while serotype 8, a non-vaccine serotype, emerged as the leading cause of IPD over the past several years (Figure 2). The incidence of IPD due to serotypes 6C, 10A, 10B, 12F, 15A, 16F, 22F, 23B, and 24F was also significantly higher by 2014/15 than would have been expected without introduction of the vaccine. The incidence of IPD due to serotype 19A increased by >3.5-fold following introduction of PCV7 but returned to levels similar to or below baseline following the introduction of PCV13 (Figure 2)

**Table 1.** Changes in IPD incidence caused by PCV7/13 serotypes and non-vaccine serotypes by age presence of comorbidity for 2014/15 compared with values expected in the absence of vaccination.

| Age group | Comorbidities<sup>1</sup> | % of IPD preventable by PCV7/13<sup>2</sup> | % Change IPD by PCV13 serotypes (95% CIs)<sup>3</sup> | % Change IPD by non-vaccine serotypes (95% CIs)<sup>3</sup> | Net % Change (95% CIs) |
| --- | --- | --- | --- | --- | --- |
| <5 years | N | 92% | -94% (-97%, -91%) | 178% (60%, 331%) | -72% (-80%, -59%) |
|  | Y | 83% | -92% (-96%, -84%) | 99% (19%, 241%) | -64% (-77%, -44%) |
| 5-17 years | N | 81% | -73% (-84%, -51%) | 57% (-21%, 151%) | -50% (-67%, -27%) |
|  | Y | 74% | -81% (-90%, -66%) | 91% (13%, 223%) | -49% (-67%, -24%) |
| 18-39 years | N | 74% | -72% (-82%, -58%) | 75% (7%, 167%) | -44% (-59%, -25%) |
|  | Y | 86% | -83% (-90%, -72%) | 51% (-2%, 126%) | -50% (-64%, -31%) |
| 40-64 years | N | 82% | -70% (-78%, -59%) | 103% (45%, 178%) | -26% (-40%, -8%) |
|  | Y | 75% | -81% (-86%, -73%) | 48% (14%, 93%) | -37% (-49%, -23%) |
| ≥65 years | N | 66% | -79% (-85%, -72%) | 60% (19%, 110%) | -44% (-54%, -31%) |
|  | Y | 70% | -78% (-83%, -71%) | 94% (53%, 144%) | -26% (-38%, -12%) |

<sup>1</sup>Any comorbidity, as defined by ACIP according to ICD10 codes registered during and hospitalization prior to the hospitalization for IPD, N= not known, Y= any comorbidity. <sup>2</sup>The % of IPD preventable by PCV7/13 is calculated using the counterfactual estimates from the model for 2014/15 by dividing the values for PCV7/13 serotypes by the overall counterfactual predicted number of cases. <sup>3</sup>The percent (%) change for both PCV13 serotypes and non-PCV13 serotypes was calculated by comparing the fitted values for 2014/15 with the counterfactual estimates from the model for 2014/15, which give an estimate for how many cases would have occurred in the absence of vaccine introduction and expressed as the estimated with 95% credible intervals (95% CIs).

**Figure 2:**
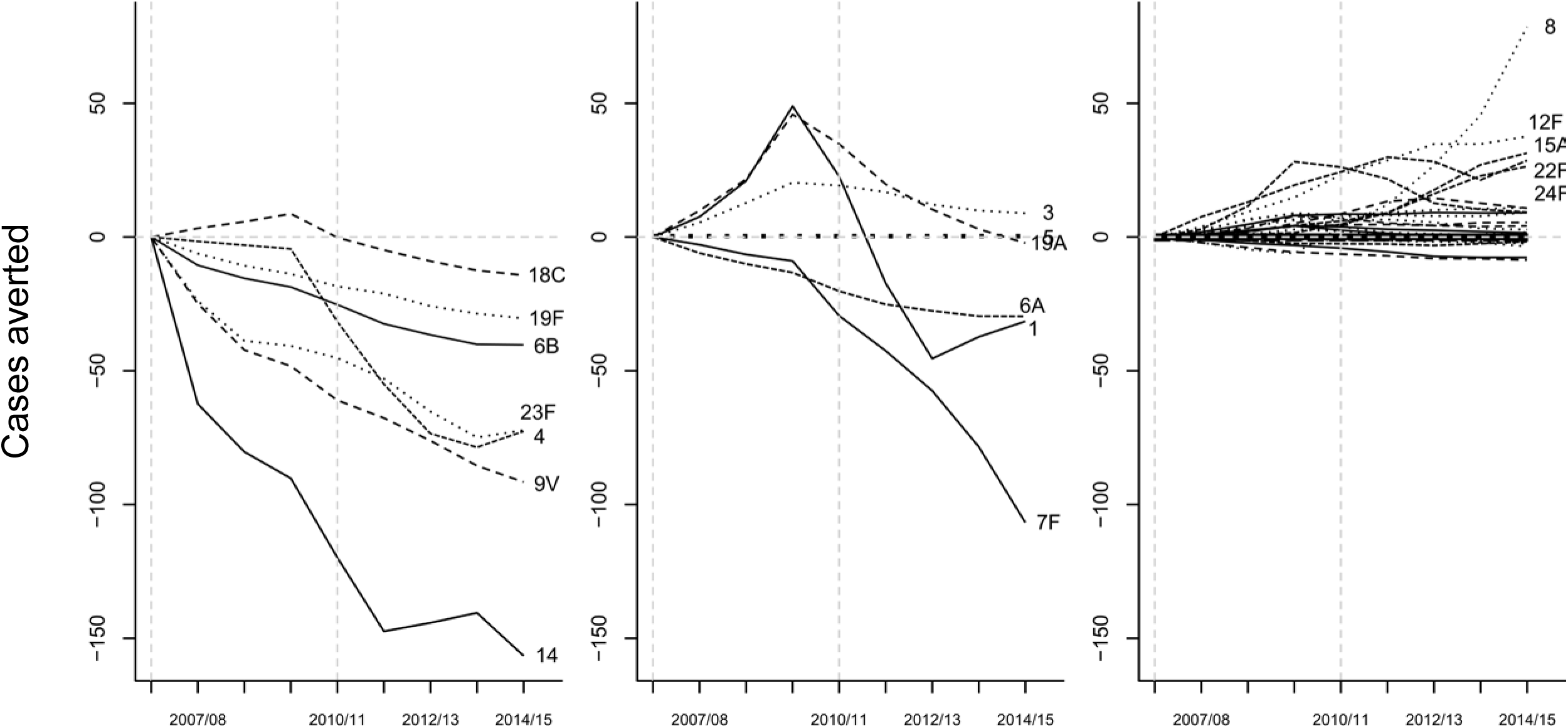
Estimated cases of invasive pneumococcal disease in all age groups prevented by vaccination (negative values) or gained through serotype replacement (positive values) in each year since the introduction of PCV7/13. Stratified by PCV7-targeted serotypes, PCV13-additional serotypes, and non-vaccine serotypes. The dashed vertical lines indicate the introduction of PCV7 and PCV13.

### The importance of serotype replacement varies by age and comorbidity level

By 2014/15 (4 years post-PCV13), there were substantial net declines in the incidence of IPD in all age groups. Incidence declined by 70% (95% Credible Intervals (CI) 57%-78%) among children <5 years of age during this time period. In contrast, the incidence of IPD among adults ≥65 years of age with comorbidities declined by just 26% (95% CI 13%-37%) (Table 1, Figure 3). Serotype replacement offset the declines in the incidence of IPD due to vaccine-targeted serotypes, but the net effect of these increases varied by age group and those with or without comorbidity. Among children <5 years of age, serotype replacement offset a small amount of the decline in IPD due to vaccine serotypes—without replacement we would have achieved an 86% reduction in IPD, and with replacement we achieved a 72% decline (Table 1, Figure 3). In contrast, among adults ≥65 with any comorbidities, serotype replacement offset more than half of the reduction in IPD caused by vaccine serotypes by the end of the study period (i.e., we would have achieved a 54% reduction In IPD without serotype replacement but instead achieved a 26% reduction) (Table 1, Figure 3).

**Figure 3:**
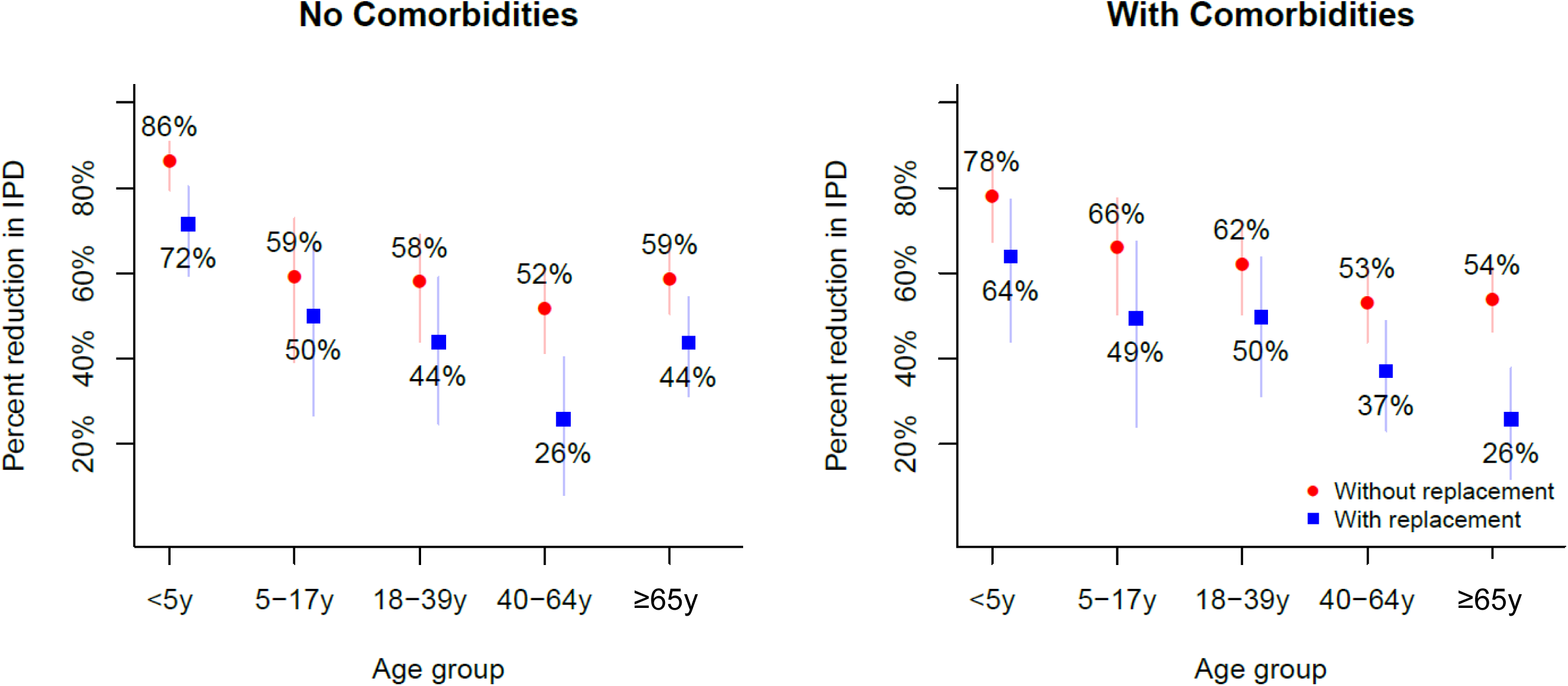
Reduction in invasive pneumococcal disease and the effect of serotype replacement, by age group and comorbidity status. The blue dots indicate the actual estimated reduction in invasive pneumococcal disease that results from both declines in vaccine-targeted serotypes and increases in non-vaccine serotypes. The red dots indicate the reduction in the incidence of invasive pneumococcal disease that would have been expected if there had been no serotype replacement. The distance between the blue and red dots indicates how much serotype replacement reduces the benefit of the vaccine. Bars indicate 95% credible intervals.

The importance of serotype replacement was particularly notable following the introduction of PCV7 and prior to the introduction of PCV13. During this time, the IPD incidence among children <5 years of age declined substantially despite replacement. However, among older children and adults, the increase in IPD due to non-vaccine serotypes completely offset declines in IPD due to vaccine-targeted serotyped (i.e., there was no net change in the overall incidence of IPD) (Figure S1).

### Differences in the net effect of replacement on IPD are due to variations in baseline incidence

We next evaluated why the net effect of the vaccine on rates of IPD differed by age group and comorbidity level. Such differences between groups could be due to variations in the pre-vaccine incidence of IPD due to non-vaccine serotypes relative to vaccine serotypes or to the magnitude of the relative increases or decreases. As could be expected, there was a strong association between the proportion of cases of IPD that were due to PCV7/13-targeted serotypes in a given age or comorbidity group prior to the introduction of PCV7/13 and the magnitude of the net decline in the incidence of IPD in that group (Table 1, Figure S3, Figure S4).

The relative increases in the incidence of IPD due to non-vaccine serotypes were notably similar across age groups (with and without comorbidity) (Table 1, Figure 4). There was modest variability between age/comorbidity groups in the strength of the relative decline in the incidence of IPD due by PCV-targeted serotypes. The declines were greatest (94%) in children <5 years of age (the only age group in which a high proportion of individuals received the vaccine) (Table 1) and with smaller, but substantial (72-83%), declines in other age groups with or without comorbidity (Table 1). By 2014/15 (7 years after the introduction of PCV7 and 4 years after the introduction of PCV13), IPD caused by PCV7-targeted serotypes had been nearly eliminated as causes of disease. The exception was IPD due to serotype 19F, which declined by 75%.

**Figure 4:**
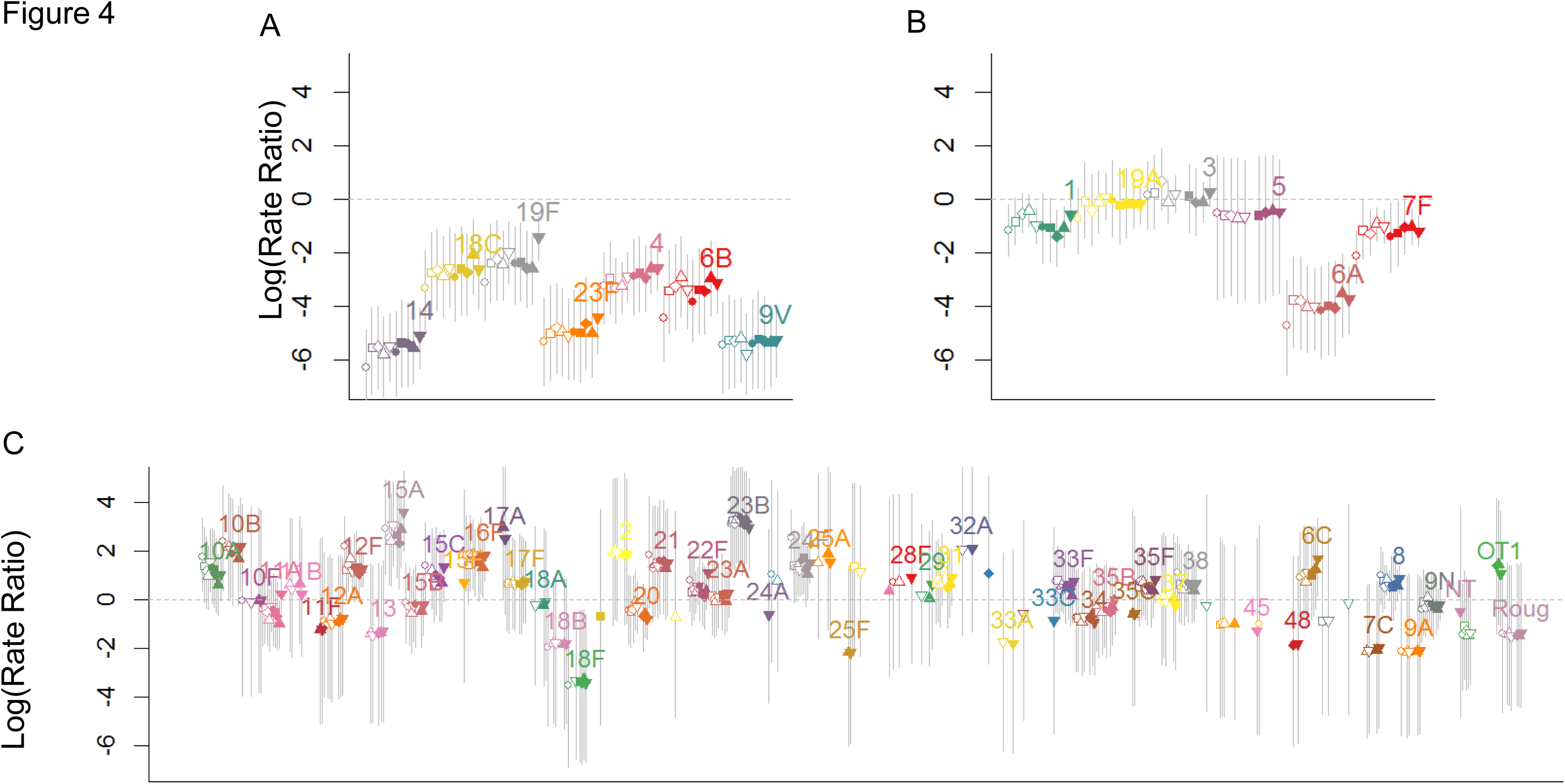
Estimated vaccine-associated change for each serotype and risk group for 2014/15. Log (rate ratios) +/- 95% credible intervals. The dots are ordered by age group (<5, 5-17, 18-39, 4064, ≥65y) and by comorbidity (open symbols: no comorbidity, closed circles: at least one comorbidity). (A): PCV7 serotypes (B) PCV13-unique serotypes (C) Non-PCV13 serotypes.

## DISCUSSION

There were clear and substantial declines in the incidence of IPD due to vaccine-targeted serotypes following the introduction of PCVs in children, as well as smaller but substantial increases in the incidence of IPD due to non-vaccine serotypes. The net effect of these changes varied somewhat by age and comorbidity status. Increases in IPD due to non-vaccine serotypes offset more of the decline in IPD due to vaccine serotypes following the introduction of PCV7 than PCV13. By using a modeling strategy that controlled for the natural epidemic cycles of IPD by the individual serotypes, it was possible to obtain credible estimates for serotype-specific changes associated with the introduction of PCVs in different groups of patients, despite natural fluctuations in the serotype-specific incidence of IPD.

The impact of PCVs against IPD in children has been well-established in other studies.^1-4^ However, there is substantial variability in estimates of the indirect benefits of the vaccine for IPD in adults and in estimates of the importance of serotype replacement. For instance, a meta-analysis by Feikin et al.^1^ found that in some countries (e.g., the US), there were significant declines in the overall incidence of IPD despite replacement. However, on average across the included countries, increases in the incidence of IPD due to non-PCV7 serotypes among adults completely offset declines in IPD due to PCV7-targeted serotypes. Our results could help to explain these patterns. We found that among adults aged 40-64 years with known comorbidities, IPD serotype replacement following the introduction of PCV7 offset declines in IPD by vaccine-targeted serotypes. Among all age groups, the relative increases in the incidence of IPD due to the non-vaccine serotypes were similar across age and comorbidity levels, and the relative declines in IPD due to vaccine serotypes were also similar. Therefore, the differences in the net effect of the vaccine between groups can be explained by serotype patterns in the period before PCV7 was introduced. In particular the balance of vaccine serotypes to non-vaccine serotypes during the pre-PCV period is important. Such variations in IPD serotype distribution could occur due to differential invasiveness of some serotypes in children and adults or due to variations in exposure between populations ^21^. In groups in which a large fraction of IPD in the pre-PCV period was due to non-vaccine serotypes, serotype replacement offset a larger fraction of the decline in IPD.

We found that the relative decline in the incidence of IPD due to PCV-targeted serotypes was greatest in children <5 years of age. This pattern is consistent with these children benefiting from both direct protection by the vaccines against invasive infections and from the decline in exposure to the vaccine-targeted serotypes that results from PCVs effect on reducing carriage of vaccine serotypes. Because effectiveness of the vaccine against targeted serotypes is greater for IPD than for carriage,^22,23^ we would expect that directly vaccinated individuals would have larger declines in the incidence of IPD due to PCV-targeted serotypes. The similar increases in the rate ratios for non-vaccine serotype IPD across all age and comorbidity groups is also consistent with the notion that all groups share a similar source of exposure (i.e., children who carry the bacteria in the nasopharynx). As a result, changes in carriage would be expected to result in proportional changes in the incidence of IPD, assuming that the serotype-specific virulence of the pneumococci does not change and that the susceptibility of the host population does not change.

Our study has strengths as well as limitations. A strength is that we used data from a comprehensive national surveillance system with a long baseline period, which allows us to estimate secular trends and harmonic variations. Additionally, we used a hierarchical modeling approach that reduces the effect of random variation in the disaggregated data and that allowed us to obtain estimates of the effect of PCVs on the incidence of IPD for different subgroups. A limitation is that we defined comorbidity status based on classification of comorbidities from a national hospitalization register, which might result in misclassification and underestimation of individuals presenting with less severe comorbid disease, particularly if these patients had not been hospitalized prior to the IPD episode. Other factors that are well known to affect susceptibility to IPD and possibly to IPD due to specific serotypes, like smoking, crowding, socioeconomic status and previous vaccination with any pneumococcal vaccine were not included in the analysis. We also assumed that linear trends continue indefinitely into the post-vaccine period. This could result in over- or under-estimating declines depending on the direction of the secular trends. Non-parametric or time series forecasting methods could help to mitigate this issue.

In conclusion, using a model that accounts for natural epidemic variations, we have demonstrated important increases in the number of cases of IPD caused by non-vaccine serotypes that accompanied declines in cases caused by vaccine-targeted serotypes following the introduction of PCV7 and PCV13 in children. The importance of these changes varied by age and comorbidity group, particularly during the period after introduction of PCV7 and before introduction of PCV13. These detailed findings will help to predict the impact of serotype replacement on IPD in different risk group following the introduction of next-generation PCVs and could assist the selection of serotypes that should be included in new PCVs for use in adults.

---

**Figure S1:**
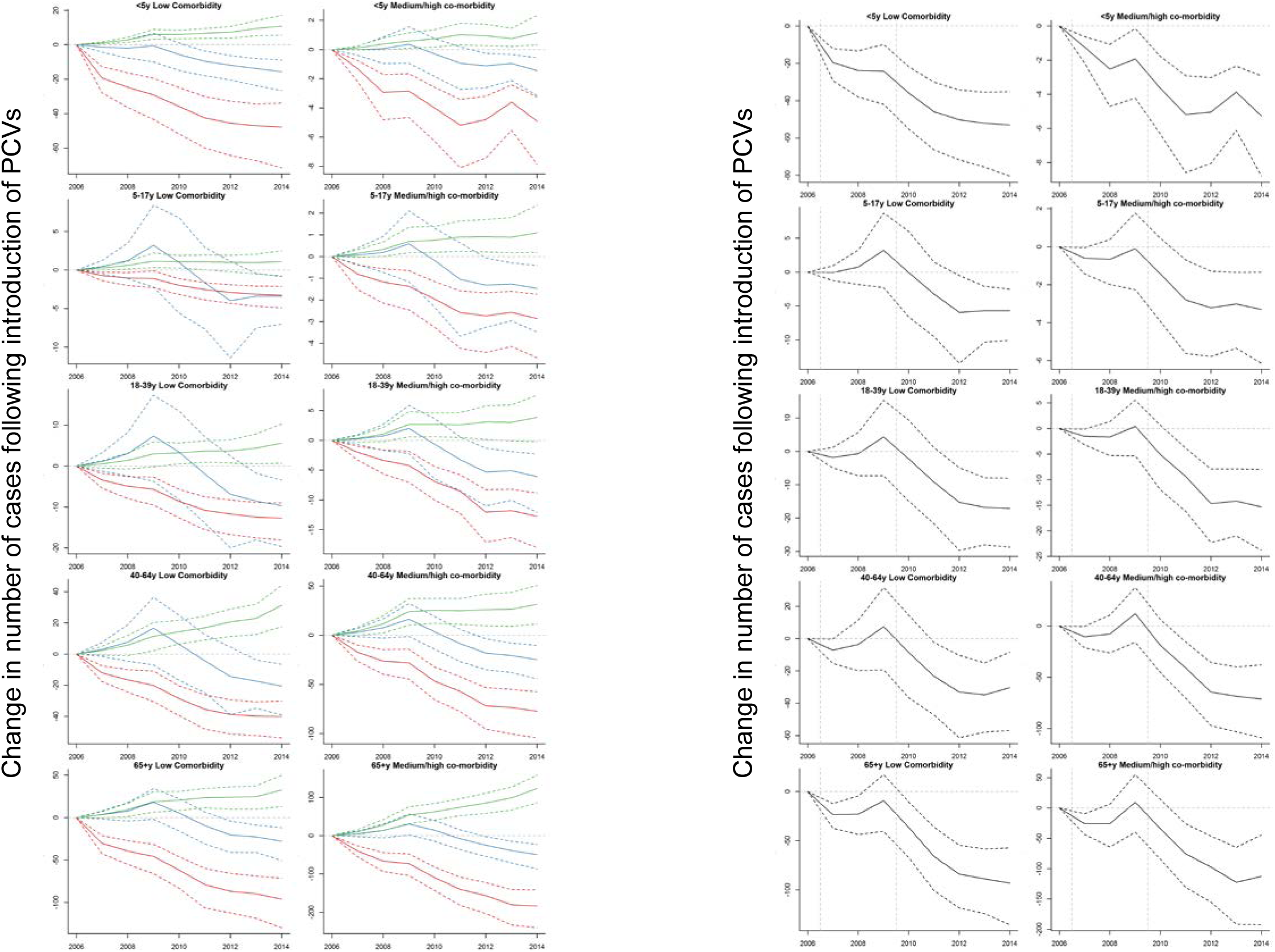
Estimates of cases averted in each year broken down by PCV7 serotypes (red), PCV13-specific serotypes (blue) and non-PCV serotypes (green). The right panel shows the net change, which is a sum of the three lines on the left. Cases prevented+/- 95% credible interval. Negative numbers indicate incidence is lower than expected following vaccine introduction, positive numbers indicate an increase.

**Figure S2:**
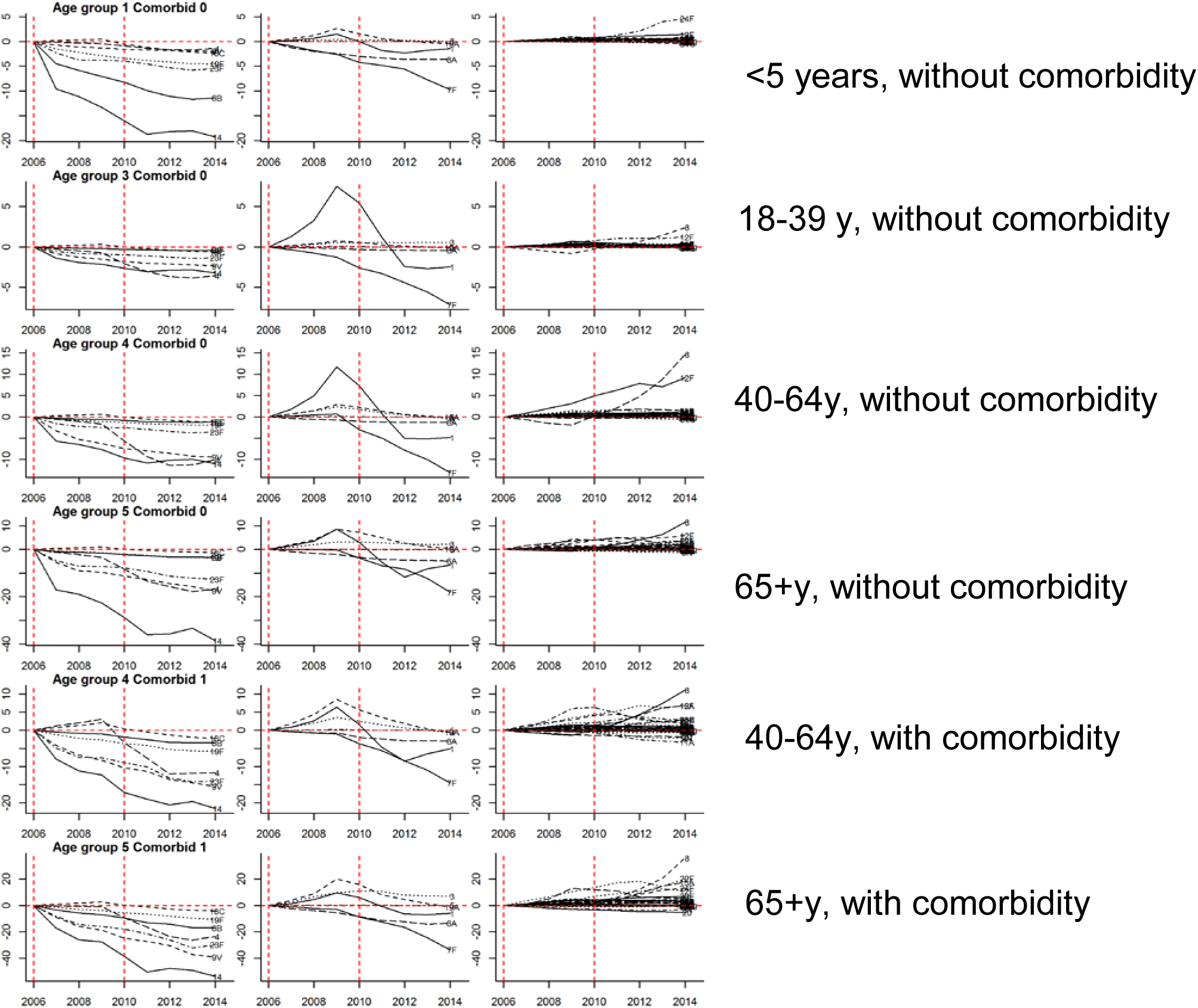
Cases averted for each serotype in selected risk groups in each year compared with the expected value. The red lines indicate the introduction of PCV7 and PCV13, respectively.

**Figure S3:**
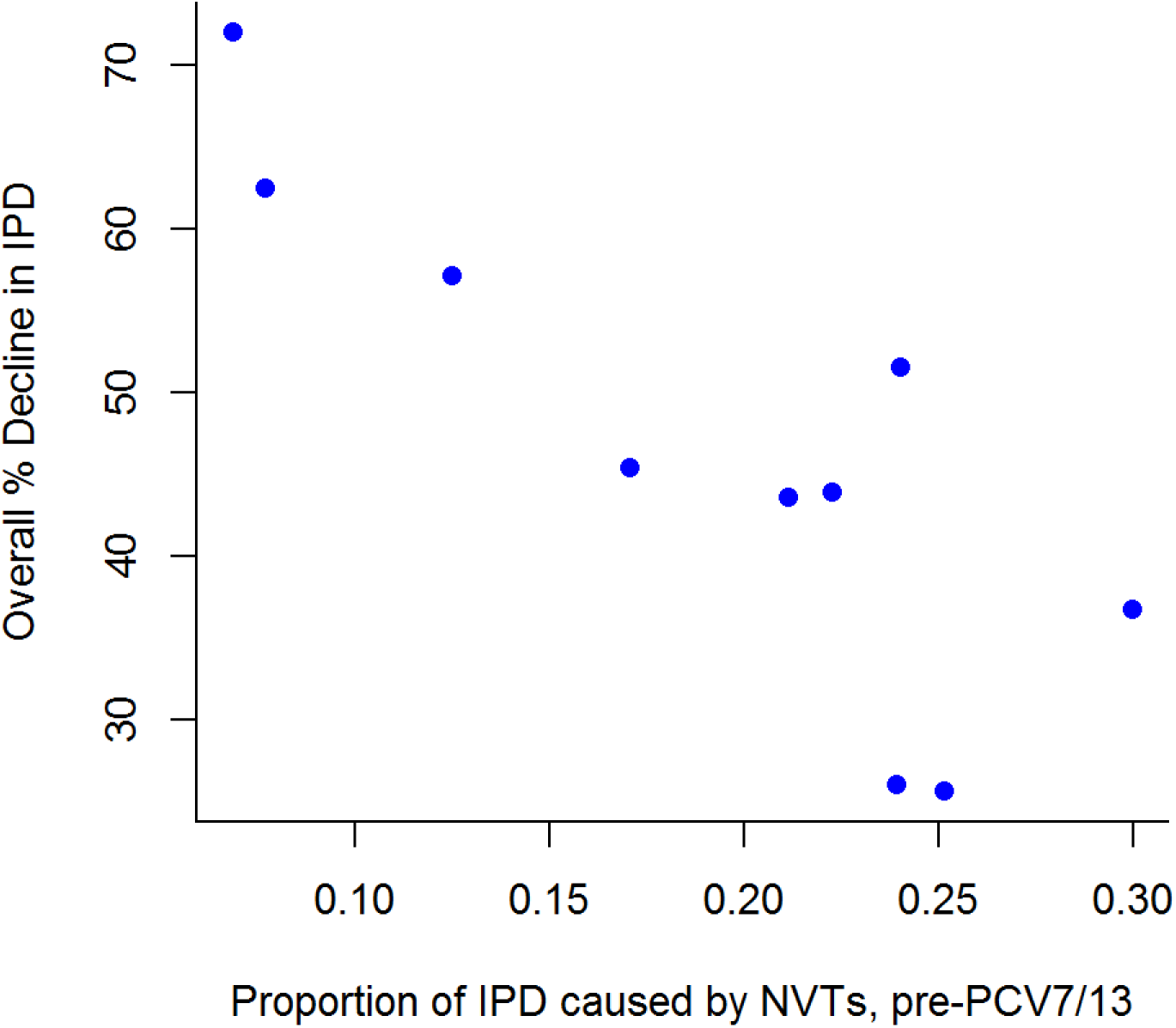
Association between the fraction of cases caused by PCV13 serotypes in the 3 years prior to introduction of PCV7 and the net decline in IPD incidence by 2014/15.

**Figure S4:**
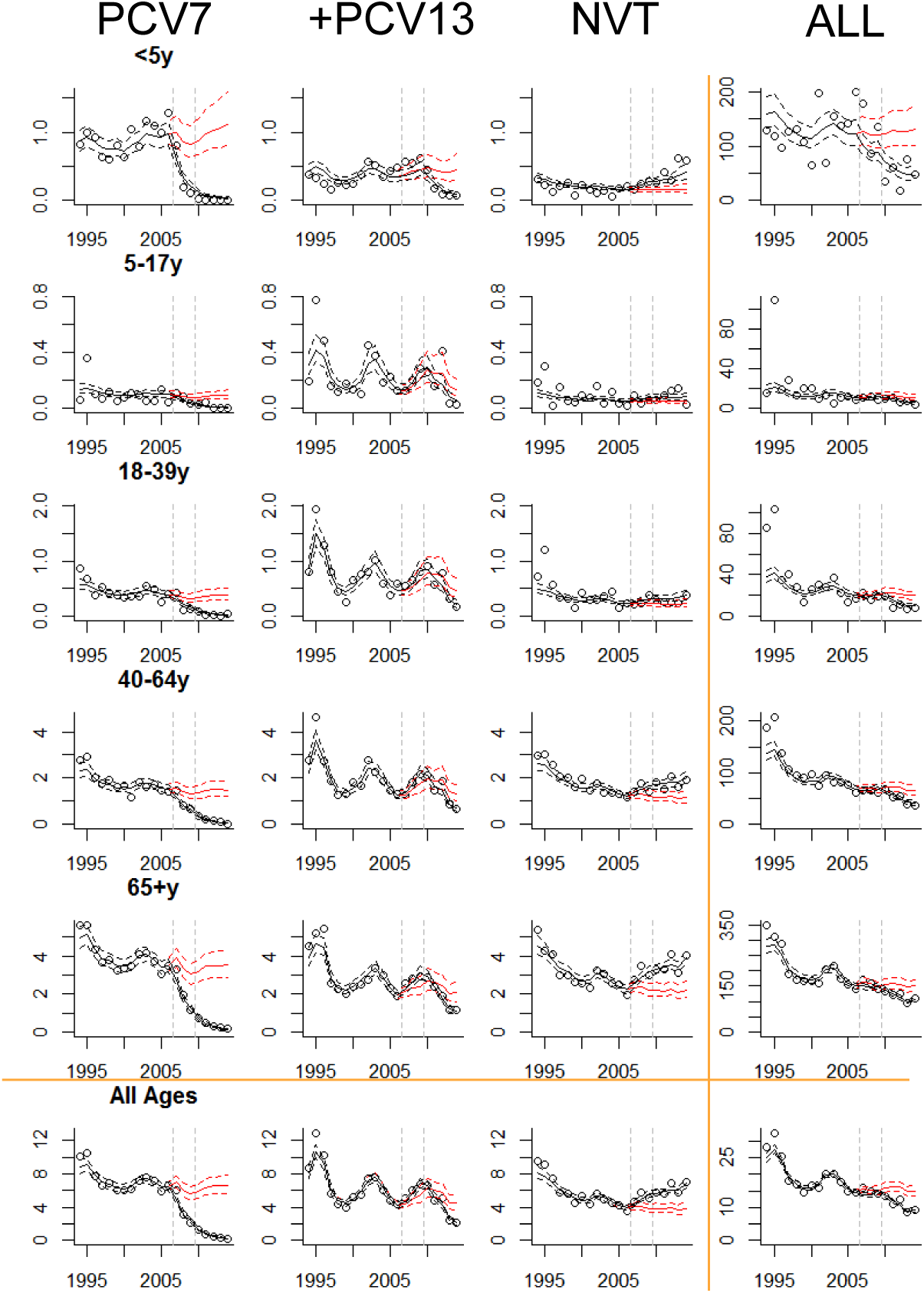
Changes in the incidence of invasive pneumococcal disease (cases/100,000) from 1994/95 to 2014/15 for serotypes targeted by PCV7, for the 6 additional serotypes in PCV13, for non-PCV13 serotypes, and for all cases combined. Different age groups are shown, as well as all age groups combined. Incidence is standardized to the age and comorbidity distribution for 2000/01, chosen arbitrarily. The dots represent observed values, the black line+/- 95% credible intervals indicates the model-fitted values, and the red line +/- 95% credible intervals represents the estimate of what the incidence would have been in the absence of vaccine.

**Figure S5:** Harmonic components from the regression model estimated for each serotype. Mean +/- 95% credible intervals.

**Figure S6:** Observed vs expected incidence of IPD for each serotype, age groups, comorbidity-level stratification. Dots indicate observed values, black line indicates the fitted value, red line +/- 95% CI indicates expected incidence in the absence of vaccine.

## SUPPLEMENTARY METHODS

We modeled the incidence of IPD at each time point and in each strata (5 age groups, with or without comorbidity) while controlling for secular trends as well as repeating epidemic cycles. The number of IPD cases caused by serotype s in risk group g at time t:

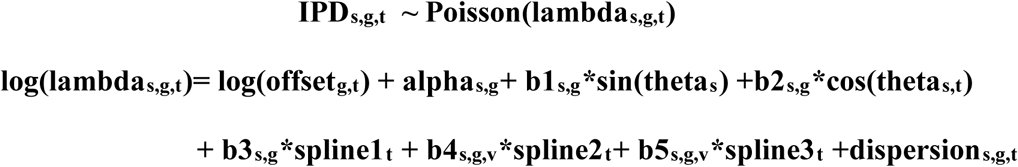

where theta_s_=2*pi*t/periods. The assumption for thetas is that the epidemic period for IPD caused by a given serotype was consistent for all risk groups, which would be expected if harmonics are driven by carriage in children ^13,24^. **spline1t, spline2t, spline3t** are modeled using B-splines, with knots at 2006/07 and 2009/10 (last time points prior to expect vaccine effect). This captures changes associated with vaccine introduction as well as secular trends that were occurring prior to the introduction of PCV7. The assumption when calculating the vaccine effect is that any trends in the pre-vaccine period continues into the post-vaccine period. In the hierarchical structure of the model, serotypes were classified (v) as PCV7 serotypes, PCV13 unique serotypes and non-vaccine serotypes. Because the data in any particular strata (serotype/comorbidity level/age-group) are sparse, we used a hierarchical model that borrows information between groups and allows for more reliable estimates than if each strata were considered separately.

Briefly, the parameters for the slopes and intercepts for each serotype and risk-group were centered around the overall serotype-level parameter. Individual slopes for changes associated with the introduction of PCV7 and PCV13 (**b3**_**s,g**_, **b4**_**s,g,v**_, **b5**_**s,g,v**_) for each serotype and risk group were centered around the overall serotype-level slope. **b4**_**s,g,v**_, **b5**_**s,g,v**_ were in turn centered around the average slope for the corresponding grouping of PCV7 serotypes, PCV13-unique serotypes, or non-PCV serotypes. This hierarchical structure allows us to appropriately quantify the variance for different levels of aggregation and “shrink” estimates towards higher level group means based on the information contained in the data. The hyperprior distributions are described in detail below.

The estimated changes associated with introduction of PCVs are calculated by estimating lambda_s,g,t_ as well as no_PCV_lambda_s,g,t_ (using the model equation above but holding b4_s,g,v_ and b5_s,g,v_ to 0 and assuming the pre-vaccine trend continued into the post-vaccine period). Cases prevented and rate ratios (RR) were then calculated as the difference or ratio between lambda_s,g,t_ and no_PCV_lambda_s,g,t_.

The models were fit using JAGS in R statistical software ^19,20^. We initialized two independent Markov chains and collected 10,000 posterior samples from each chain after a burn-in period of 5000 iterations. The 95% credible intervals (95% CI) were calculated from the 2.5^th^ and 97.5^th^ percentiles of the collected posterior samples. Convergence was assessed using Geweke’s diagnostic.^25^ Alternative models that excluded the pre-vaccine trends and that allowed the variance on the dispersion parameter to vary for each time series were also considered. The models were compared using the Deviance Information Criterion (DIC),^26^ and the final selected model for making inference had the lowest DIC value indicating an improved balance of model fit and complexity compared to other models.

The prior and hyperprior distributions for the model were specified as follows. Uniform(0,100) prior distributions were specified for all standard deviation parameters.

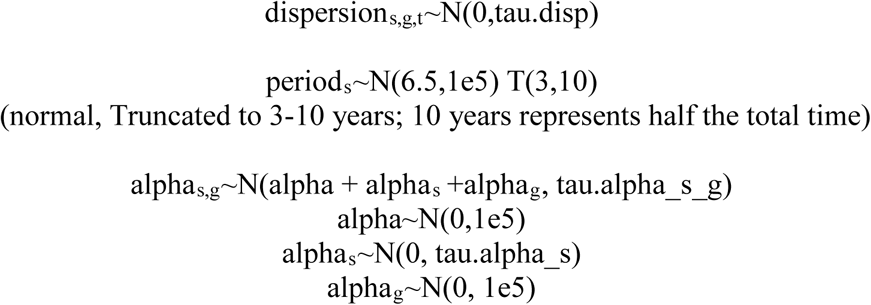

The estimate for the harmonics varies by serotype and is “shrunk” towards 0, indicating that most serotypes will have little harmonic variation

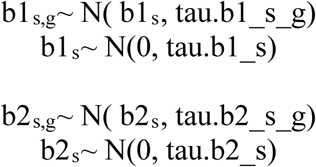

Secular trend for serotype s in risk group g centered around the secular trend for the serotype (10 risk groups total: 5 age groups × 2 comorbidity levels)

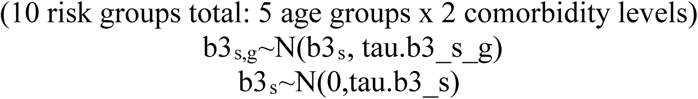

Vaccine-associated changes for serotype s and risk group g (b4_s,g,v_, b5_s,g,v_) centered around serotype changes. Serotype changes centered around average changes for the corresponding grouping (v) of serotypes. Serotypes are binned into three categories (v) based on whether they are targeted by PCV7, PCV13, or neither.

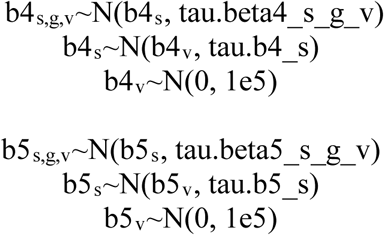

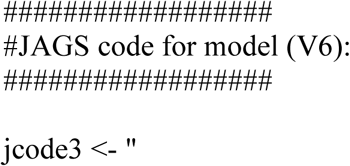

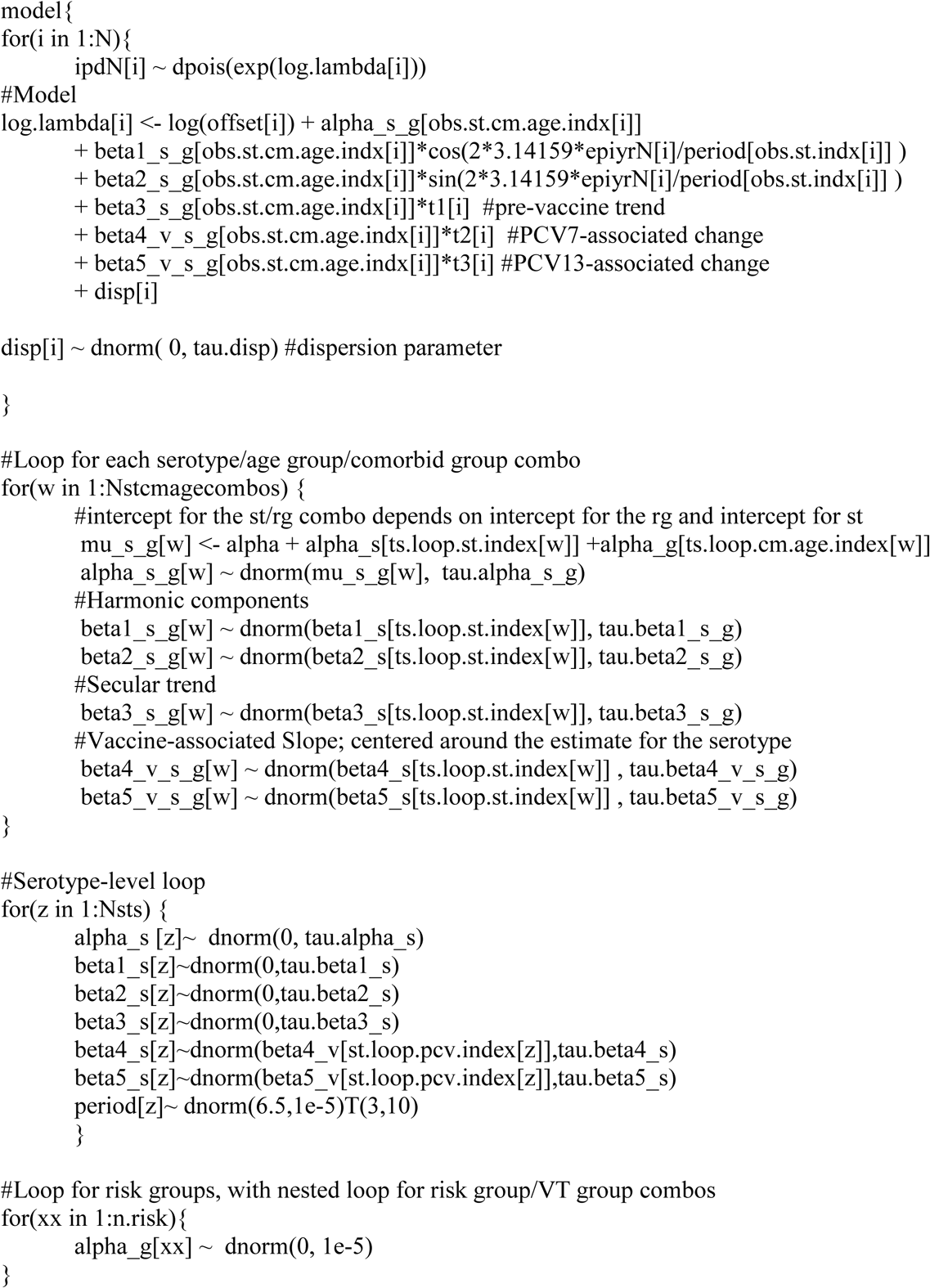

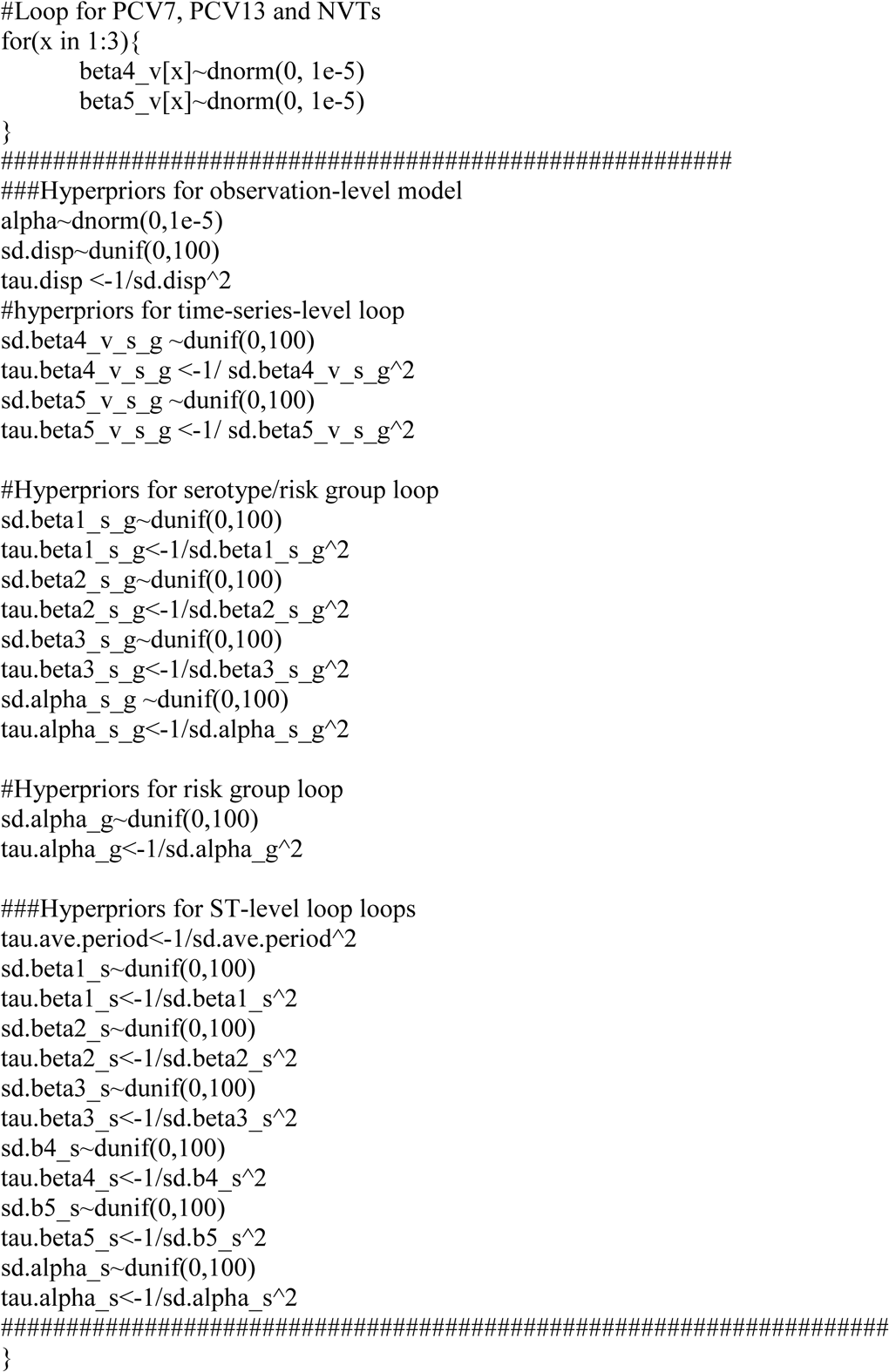

ICD10 Codes (Danish edition) used to define comorbid conditions

## Chronic Heart diseases

DI099, DI110, DI21, DI22, DI23, DI24, DI25, DI130, DI132, DI255, DI420, DI425, DI426, DI427, DI428, DI429, DI43, DI50, DP290

## Chronic Pulmonary Diseases

DI278, DI279, DJ40, DJ41, DJ42, DJ43, DJ44, DJ45, DJ46, DJ47, DJ60, DJ61, DJ62, DJ63, DJ64, DJ65, DJ66, DJ67, DJ684, DJ701, DJ703, DJ841, DJ920, DJ961, DJ982, DJ983, DE84, DJ991

## Liver Diseases

DB18, DI85, DI864, DI982, DK70, DK711, DK713, DK714, DK715, DK717, DK72, DK73, DK74, DK760, DK762, DK763, DK764, DK765, DK766, DK767, DK768, DK769, DZ944

## Renal Failure

DI120, DI13, DN00, DN01, DN02, DN03, DN04, DN05, DN07, DN08, DN11, DN14, DN18, DN19, DN25, DN26, DN27, DQ61, DZ490, DZ49, DZ992

## Haematological malignancies

DC81, DC82, DC83, DC84, DC85, DC86, DC88, DC90, DC91, DC92, DC93, DC94, DC95, DC96, DC97

## Solid malignancies with/without metastases

DC77, DC78, DC79, DC80, DC00, DC01, DC02, DC03, DC04, DC05, DC06, DC07, DC08, DC09, DC10, DC11, DC12, DC13, DC14, DC15, DC16, DC17, DC18, DC19, DC20, DC21, DC22, DC23, DC24, DC25, DC26, DC30, DC31, DC32, DC33, DC34, DC37, DC38, DC39, DC40, DC41, DC43, DC44, DC45, DC46, DC47, DC48, DC49, DC50, DC51, DC52, DC53, DC54, DC55, DC56, DC57, DC58, DC60, DC61, DC62, DC63, DC64, DC65, DC66, DC67, DC68, DC69, DC70, DC71, DC72, DC73, DC74, DC75, DC76, DC97

## Organ transplantation

DZ94-DZ949

## Anatomic or functional asplenia

DD73, D57, D561, D58, DQ890, DQ890, DZ908

## AIDS/HIV and other immunodeficiencies

DB20, DB21, DB22, DB24, DB23, DD80, DD81, DD82, DD83, DD84, DD70, DD71, DD720

## Rheumathologic diseases

DL940, DL941, DL943, DD86, DM05, DM06, DM07, DM08, DM09, DM120, DM123, DM30, DM310, DM311, DM312, DM313, DM32, DM33, DM34, DM35, DM36, DM45, DM461, DM468, DM469

## Diabetes

DE100, DE101, DE109, DE110, DE111, DE119, DE120, DE121, DE129, DE130, DE131, DE139, DE140, DE141, DE149, DE102, DE103, DE104, DE105, DE106, DE107, DE108, DE112, DE113, DE114, DE115, DE116, DE117, DE118, DE122, DE123, DE124, DE125, DE126, DE127, DE128, DE132, DE133, DE134, DE135, DE136, DE137, DE138, DE142, DE143, DE144, DE145, DE146, DE147, DE148

## Alcohol related conditions

DF10, DK860, DZ721, DR780, DT51, DK292, DG621, DG721, DG312, DK70, DI426

## Cerebrospinal fluid leak

DG960

## Cochlear implant

DZ962

## Acknowledgements

This work was funded by NIH/NIAID grants R01AI123208 and R56AI110449 to DMW. Support was also received from the Bill and Melinda Gates Foundation (OPP1176267 and OPP1114733), the National Institute on Aging Grant #P30AG021342 (Scholar at the Claude D. Pepper Older Americans Independence Center at Yale University School of Medicine) and from grants # UL1TR001863, UL1 TR000142 and KL2 TR000140 from the National Center for Advancing Translational Science (NCATS), components of the National Institutes of Health (NIH), and NIH roadmap for Medical Research. The findings and conclusions of this paper are those of the authors and do not necessarily represent the official position of the NIH or the Statens Serum Institut.

